# Melanin-concentrating hormone signaling regulates persistence and updating of reward-guided actions

**DOI:** 10.64898/2026.08.17.745216

**Authors:** Pratheba Kandasamey, Cristina Concetti, Julie Burkhardt, Eva Bracey, Denis Burdakov, Daria Peleg-Raibstein

## Abstract

Flexible behavior requires persistence when a strategy remains effective and rapid updating when its consequences change. Melanin-concentrating hormone (MCH) neurons in the lateral hypothalamus regulate feeding, reward and memory, but their contribution to reward-guided flexibility is unknown. Here, mice learned an action–outcome contingency in a T-maze and then adapted when the reward location switched. MCHR1 antagonism throughout learning did not measurably alter Initial Learning but reduced perseverative choices and accelerated behavioral adaptation after the contingency switched. Endogenous LH-MCH activity was strongest while contingencies were being established or revised, declined as performance stabilized, distinguished rewarded from unrewarded outcomes, and reflected current outcomes in the context of recent and accumulated experience. MCHR1 blockade altered prelimbic responses to successful outcomes in both phases and opposed their progressive weakening during Rule Switch. Together, these findings identify MCH signaling as a regulator of how strongly past reward continues to guide behavior when contingencies change and reveal accompanying changes in prefrontal processing of successful outcomes.

## Introduction

Adaptive behavior requires more than learning which actions lead to reward. When action–outcome contingencies change, previously successful responses must be suppressed and behavior redirected toward the currently valid alternative. This capacity is central to cognitive flexibility and depends on integrating recent outcomes with ongoing task demands to determine whether the current response policy should be maintained or revised. Failures of this process contribute to behavioral rigidity and maladaptive persistence, features relevant to compulsive reward seeking and other disorders characterized by inflexible behavior ^1–11^.

The medial prefrontal cortex (mPFC) is strongly implicated in this form of adaptive control. Across rodent and primate studies, mPFC and adjacent cingulate regions encode recent outcomes, action history, conflict, and task context in ways that bias behavior toward the currently adaptive response while suppressing responses that are no longer appropriate ^1–10,12–18^ . Within this framework, mPFC does not simply represent reward or movement, but contributes to selecting the response policy that best matches current contingencies ^8–10,12,15–18^. Prelimbic population activity is well positioned to reflect the integration of task, outcome, and rule information during adaptive response selection. ^1–3,8–10^. What remains less clear is which neuromodulatory systems regulate these cortical computations when action–outcome contingencies must be acquired and then revised. Recent work further showed that anterior cingulate populations retain information about previous choices and outcomes that is separable from ongoing posture and movement, suggesting that trial-history monitoring is a fundamental component of adaptive control ^10^.

Melanin-concentrating hormone neurons in the lateral hypothalamus have classically been studied in feeding, energy balance, and food reward. MCH neurons respond to food-predictive cues and consumption, and recent work has shown that their activity interacts with accumbens dopamine during food-motivated Pavlovian learning ^19–21^. Beyond reward processing, MCH neurons have been implicated in object memorization, REM-associated forgetting, fear extinction, and the regulation of impulsive behavior ^22–27^. These findings indicate that the MCH system can influence the formation, persistence, and modification of learned representations. An anatomical substrate for interaction with prefrontal circuits is also present: MCH-immunoreactive fibers extend throughout neocortical regions, including frontal cortex ^28^, whereas MCHR1 mRNA and protein are broadly expressed in cerebral cortex ^29^ and have been localized to cortical pyramidal neurons ^30^. Consistent with this possibility, local MCHR1 blockade within prelimbic mPFC facilitates the updating of learned avoidance ^31^. Together, this work establishes a role for MCH beyond homeostatic control, but leaves unresolved how endogenous LH-MCH activity evolves during appetitive instrumental updating, whether it carries recent-outcome information, and how MCHR1 signaling relates to prefrontal outcome activity in this context.

Here, we asked how endogenous LH-MCH activity evolves as an appetitive action–outcome contingency is first acquired and then changed; whether it incorporates recent outcome history; and whether MCHR1 antagonism alters behavioral updating and outcome-dependent prelimbic mPFC activity. We combined systemic MCHR1 antagonism with fiber photometry recordings of LH-MCH neurons and CaMKIIα-targeted prelimbic populations in a two-phase appetitive T-maze. This approach allowed us to test whether a hypothalamic system classically associated with feeding and reward also contributes to the balance between maintaining and revising learned reward-guided actions as contingencies change.

## Results

### Endogenous LH-MCH activity is recruited during contingency learning and re-engaged during Rule Switch

To examine how endogenous MCH activity changes as reward contingencies are acquired and revised, mice were trained in a two-phase appetitive T-maze. During Initial Learning, one goal arm contained a food reward and the opposite arm contained an empty cup. After animals reached criterion, the reward location was reversed during Rule Switch. We expressed GCaMP6s in lateral hypothalamic MCH neurons and recorded fiber-photometry signals aligned to four task epochs: the early inter-trial interval, late inter-trial interval, trial start, and cup entry (Fig. 1a–c). Signals were examined separately for correct and incorrect choices during the first, second, and final session of each phase.

**Figure 1.**
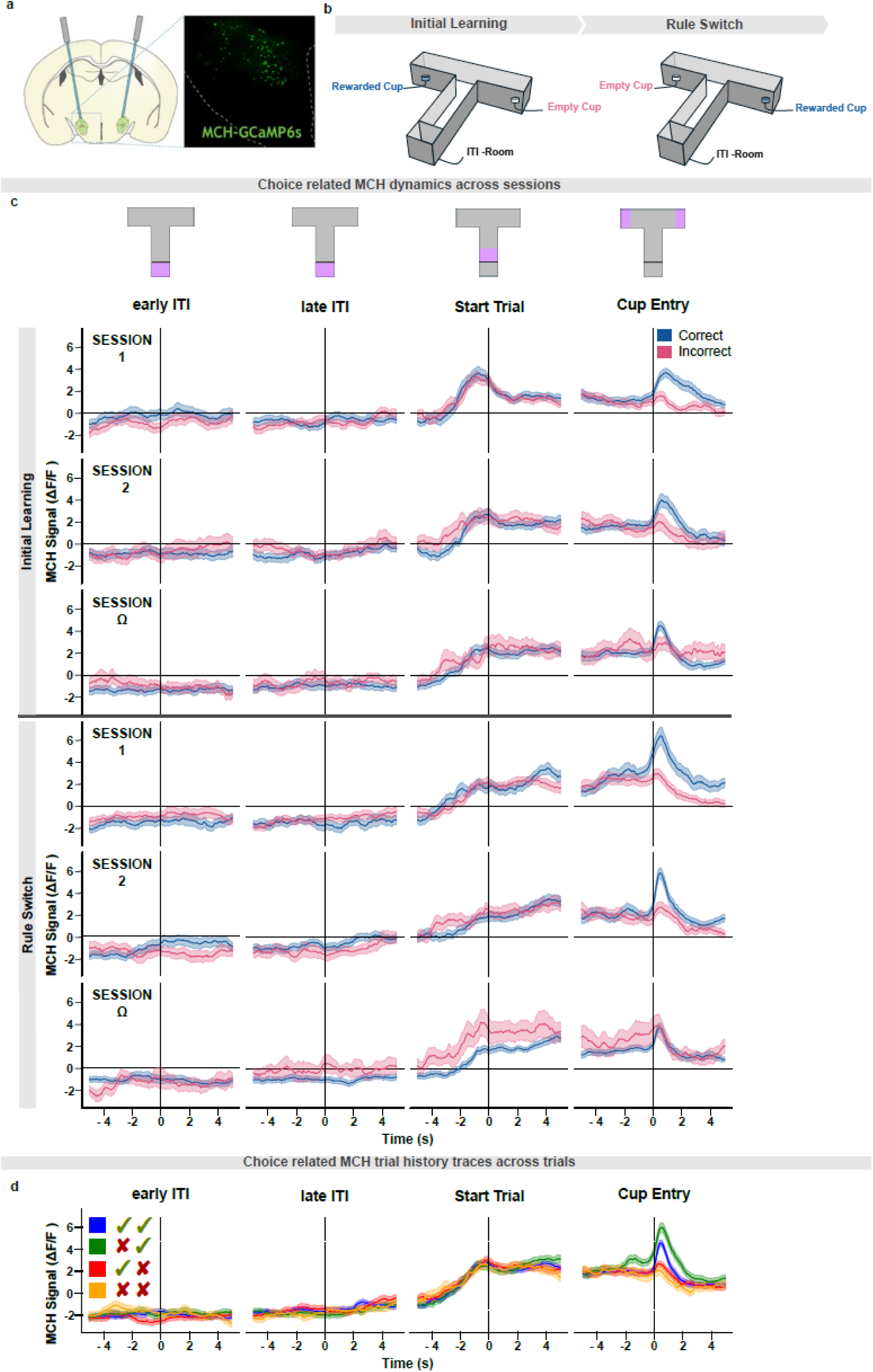
Endogenous LH-MCH activity is recruited during contingency learning and re-engaged during Rule Switch. (a) Schematic of GCaMP6s expression in lateral hypothalamic MCH neurons and optical fiber placement for fiber-photometry recording. (b) Two-phase appetitive T-maze task. During Initial Learning, one arm contained the rewarded cup and the opposite arm an empty cup. After criterion was reached, the rewarded arm was reversed during Rule Switch. (c) Trial-averaged LH-MCH calcium signals aligned to the early inter-trial interval (early ITI), late inter-trial interval (late ITI), trial start, and cup entry, shown separately for correct and incorrect trials in the first (S1), second (S2), and final (SΩ) session of each phase. (d) Descriptive one-back trial-history organization of LH-MCH traces. Trials are grouped as correct→correct (✓✓), incorrect→correct (✗✓), correct→incorrect (✓✗), and incorrect→incorrect (✗✗), providing a qualitative view of how LH-MCH activity varies with the immediately preceding and current outcomes. Signals are shown as mean ± SEM. n = 9 mice (2 male, 7 female).

Across both phases, the clearest phasic modulation occurred at cup entry. Both correct and incorrect trials were accompanied by positive cup-entry transients, but the response was larger following correct, rewarded choices. The learning-stage dependence was also most apparent on correct trials: the cup-entry response was strongest early in Initial Learning, declined as the contingency was acquired, re-emerged at the start of Rule Switch, and decreased again as animals learned the switched contingency. Incorrect-trial responses varied across sessions but did not show the same consistent trajectory. Thus, the learning-stage dependence of LH-MCH activity was most evident following rewarded outcomes and was unlikely to reflect a fixed response to food consumption alone. These outcome- and learning-related effects are quantified in Fig. 2a–f.

**Figure 2.**
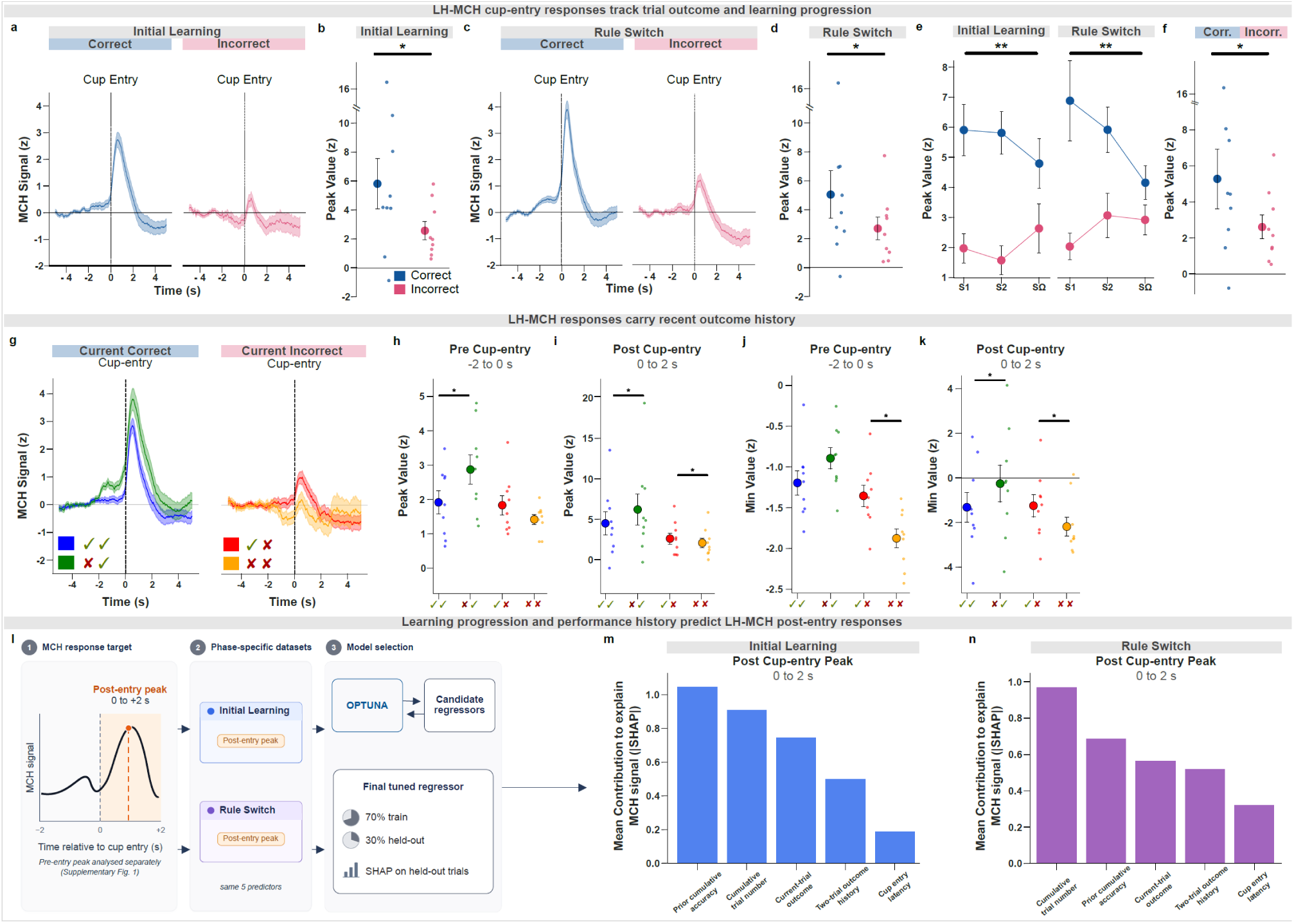
LH-MCH cup-entry responses track current outcome, learning progression, and recent outcome history. (a,c) Cup-entry-aligned LH-MCH calcium signals during Initial Learning (a) and Rule Switch (c), shown separately for correct and incorrect trials. (b,d) Subject-level quantification of post-entry peak responses during Initial Learning (b) and Rule Switch (d). (e) Peak responses across the first, second, and final sessions of each phase; statistical models included all sessions. (f) Correct versus incorrect peak responses across phases. (g) Trial-history traces grouped according to the preceding and current trial outcomes. (h,i) Peak values in the pre-entry (−2 to 0 s) and post-entry (0 to 2 s) windows; (j,k) minimum values in the corresponding windows. (l) Supervised-regression workflow for phase-specific models of the post-entry peak; pre-entry models are reported in Supplementary Fig. 1 and Supplementary Table S1. (m,n) Mean absolute SHAP feature contributions during Initial Learning (m) and Rule Switch (n); full SHAP distributions are shown in Supplementary Fig. 1. Panels b and d were analyzed using paired two-tailed Wilcoxon signed-rank tests, panel f using a paired t test, and panels h–k using two-tailed paired tests with false-discovery-rate correction. Individual points represent subject-level values. n = 9 mice (2 male, 7 female).

Adaptive updating requires animals to integrate outcomes across trials rather than evaluate each choice in isolation. Recent work showed that frontal cortical populations retain information about previous choices and outcomes that is separable from ongoing movement signals, supporting the view that trial-history monitoring is a core component of adaptive control ^10^. Motivated by this framework, we asked whether LH-MCH activity similarly reflected recent outcome context. Current-correct and current-incorrect trials were separated according to whether the immediately preceding trial had been correct or incorrect, yielding four one-back outcome sequences (Fig. 1d). Visual inspection suggested that LH-MCH responses varied across the one-back outcome sequences, particularly for current-correct trials. These patterns are shown descriptively in Fig. 1d and quantified formally in Fig. 2g–k.

### LH-MCH cup-entry responses track trial outcome, learning progression, and recent outcome history

We next quantified the cup-entry component of the LH-MCH signal. Both rewarded and unrewarded cup entries elicited positive transients, but peak responses were significantly larger on correct than incorrect trials during Initial Learning (paired Wilcoxon signed-rank test, W = 5.00, p = 0.04; Fig. 2a,b) and Rule Switch (W = 5.00, p = 0.04; Fig. 2c,d). When responses were averaged across phases, correct trials again showed larger cup-entry peaks than incorrect trials (paired t test, t(8) = 2.46, p = 0.04; Fig. 2f). Cup entry therefore elicited a common LH-MCH response under both outcomes, with an additional increase following successful reward delivery.

We next examined how correct- and incorrect-trial peak responses evolved across learning. For visualization, peak responses are shown for the first, second, and final sessions of Initial Learning and Rule Switch (Fig. 2e), whereas the mixed-effects models included data from all sessions within each phase. Outcome, Session, and their interaction were included as fixed effects, with Subject as a random intercept. Correct trials and Session 1 served as the reference levels.

During Initial Learning, the model-estimated simple effect of Outcome at Session 1 showed that incorrect-trial peaks were lower than correct-trial peaks (β = −4.02, SE = 0.86, z = −4.68, p < 0.001). The model coefficient for Session in correct trials showed that correct-trial peaks declined across sessions (β = −0.72, SE = 0.30, z = −2.38, p = 0.02). The significant Outcome × Session interaction indicated that the session-related trajectory differed between correct and incorrect trials, with a flatter trajectory for incorrect trials (β = 1.06, SE = 0.40, z = 2.62, p = 0.009; Fig. 2e). During Rule Switch, the model-estimated simple effect of Outcome at Session 1 again showed lower peaks on incorrect than correct trials (β = −4.34, SE = 0.91, z = −4.80, p < 0.001). The model-estimated Session slope for correct trials was also negative (β = −1.02, SE = 0.33, z = −3.06, p = 0.002), and the significant Outcome × Session interaction indicated a flatter trajectory for incorrect trials (β = 1.28, SE = 0.45, z = 2.85, p = 0.004; Fig. 2e).

The initially large correct–incorrect peak difference therefore decreased as each contingency was learned. Notably, the correct-trial peak increased again at the start of Rule Switch relative to the final session of Initial Learning (planned one-sided paired t test, t(6) = 2.14, p = 0.04). Together, the decline with learning and re-emergence after the reward switch indicate that outcome-related LH-MCH recruitment is strongest while the current contingency is being acquired or must be reconsidered.

We next formally quantified the one-back trial-history structure identified in the descriptive traces shown in Fig. 1d. When the current trial was correct, peak values differed according to whether the preceding trial had been correct or incorrect in both the pre-cup-entry window (pFDR = 0.01) and the post-cup-entry window (pFDR = 0.02; Fig. 2g–i). When the current trial was incorrect, the same comparison was significant only after cup entry (pFDR = 0.02) and not before cup entry (pFDR = 0.16; Fig. 2g–i). Minimum values showed a related but distinct pattern. For current-correct trials, minimum values differed after cup entry but not before cup entry (pFDR = 0.03 and pFDR = 0.08, respectively; Fig. 2j,k). For current-incorrect trials, minimum values differed in both the pre- and post-entry windows (pFDR = 0.01 and pFDR = 0.02, respectively; Fig. 2j,k). These findings show that identical current outcomes elicited different LH-MCH responses depending on the immediately preceding outcome. Recent-outcome effects were detected in both peak and minimum measures and in both pre- and post-entry windows, although the specific significant comparisons varied according to current outcome and response measure. Peak and minimum measures were analyzed separately and were not directly compared statistically.

To complement the trial-history analyses, we used supervised regression to model trial-to-trial variation in the post-entry peak separately during Initial Learning and Rule Switch (Fig. 2l). The same model framework incorporated current-trial outcome, two-trial outcome history, cumulative trial number, prior cumulative accuracy, and cup-entry latency, allowing these co-occurring behavioral variables to be evaluated jointly (Supplementary Table S2). We focused the supervised analysis on peak responses because the peak was the primary cup-entry measure used to characterize outcome- and learning-related LH-MCH recruitment (Fig. 2a–f); minimum values were retained for the targeted trial-history analysis. Model family and hyperparameters were selected using Optuna ^32^, and predictive performance was evaluated on held-out trials. Both post-entry models performed above the permutation-derived null and showed positive animal-centered R² values (Initial Learning: pooled held-out R² = 0.23, animal-centered R² = 0.10; Rule Switch: pooled held-out R² = 0.42, animal-centered R² = 0.35; Supplementary Table S1). These models were therefore retained for SHapley Additive exPlanations (SHAP) ^33^ analysis (Fig. 2m,n). By contrast, although the corresponding pre-entry models showed above-null pooled performance, their bootstrap confidence intervals included zero and animal-centered R² was approximately zero or negative; they were therefore treated as secondary analyses (Supplementary Fig. 1 and Supplementary Table S1).

In both phases, prior cumulative accuracy and cumulative trial number were the two largest contributors to model predictions, but their relative ranking differed between the phase-specific models (Fig. 2m,n). Prior cumulative accuracy ranked highest during Initial Learning, whereas cumulative trial number ranked highest during Rule Switch. Because the models were fitted separately and these predictors are correlated, this rank difference is interpreted descriptively rather than as a formal cross-phase comparison of feature importance. More generally, SHAP attribution is model-dependent and correlated predictors can share or redistribute importance; these profiles therefore describe how the fitted models used the available features rather than unique, independent or causal effects ^34^.

Taken together, the outcome, learning, trial-history, and multivariate analyses (Fig. 2a–n) show that LH-MCH cup-entry activity is organized around the demands of adaptive reward-guided behavior rather than reward receipt alone: it differentiates current outcomes, is strongest while contingencies are being established or revised, and reflects those outcomes in the context of recent trial history and accumulated experience.

### Systemic MCHR1 antagonism reduces persistence of the previously rewarded response during Rule Switch

Because endogenous LH-MCH activity was elevated while the initial action–outcome contingency was being established and again when that contingency had to be updated during Rule Switch, we administered the MCHR1 antagonist SNAP-94847 or vehicle before every behavioral session during both Initial Learning and Rule Switch (Fig. 3a). This design tested the contribution of MCHR1 signaling across the full adaptive-learning sequence rather than only during Rule Switch. Mice performed 12 trials per session and advanced when they reached the predefined criterion of at least 75% correct choices across the session and correct choices on at least 2 of the final 3 trials.

**Figure 3.**
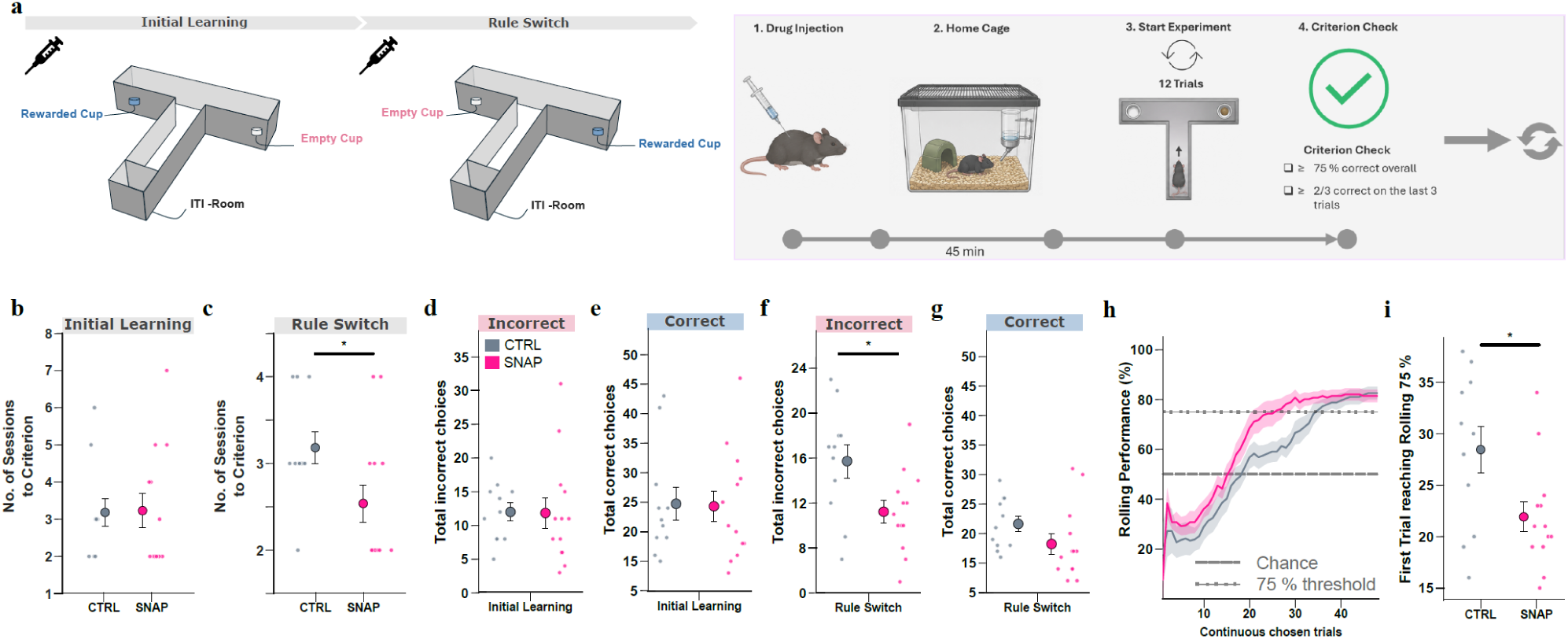
Systemic MCHR1 antagonism reduces persistence of the previously rewarded response during Rule Switch. (a) T-maze contingencies, session structure, and progression criteria. Before each session in both phases, mice received SNAP-94847 or vehicle and were returned to the home cage for 45 min. Each session consisted of 12 trials. Progression required at least 75% correct choices across the session and correct choices on at least 2 of the final 3 trials. (b,c) Sessions required to reach criterion during Initial Learning and Rule Switch. (d,e) Total incorrect and correct choices during Initial Learning. (f,g) Total incorrect and correct choices during Rule Switch. (h) Rolling performance across continuous chosen trials during Rule Switch, calculated using a 12-trial window. For visualization, each animal’s final value was carried forward after completion. Thick lines show group means and shaded regions show SEM; dashed lines mark chance performance (50%) and the 75% threshold. (i) First continuous chosen trial at which each animal reached at least 75% correct performance in the rolling window. Individual points represent mice; large symbols and error bars indicate mean ± SEM. Control, n = 11 mice (9 male, 2 female); SNAP, n = 13 mice (10 male, 3 female).

Sessions required to reach criterion were analyzed using a mixed-design ANOVA with Group as a between-subject factor and Phase as a within-subject factor. There was no significant main effect of Group (P = 0.380), main effect of Phase (P = 0.267), or Group × Phase interaction (P = 0.307; Fig. 3b,c). In planned phase-wise comparisons, Control and SNAP-treated mice did not differ during Initial Learning (P = 0.935), whereas SNAP-treated mice required fewer sessions during Rule Switch (P = 0.033). Thus, SNAP was associated with faster criterion attainment during Rule Switch, although the absence of a Group × Phase interaction does not establish that the treatment effect differed between phases.

We next examined incorrect and correct choices separately within each phase. During Initial Learning, neither total incorrect choices (Welch’s t test, t(19.25) = 0.06, P = 0.95; Fig. 3d) nor total correct choices (Welch’s t test, t(21.45) = 0.11, P = 0.91; Fig. 3e) differed between groups. During Rule Switch, SNAP-treated mice made significantly fewer incorrect choices than controls (Welch’s t test, t(18.08) = 2.50, P = 0.02; Fig. 3f), whereas total correct choices did not differ significantly between groups (Welch’s t test, t(21.36) = 1.56, P = 0.13; Fig. 3g). Thus, during Rule Switch, SNAP reduced selections of the previously rewarded, now-unrewarded arm without significantly altering the total number of correct choices. This pattern is consistent with reduced persistence of the outdated response strategy.

A trial-resolved analysis yielded the same conclusion. Rolling performance across continuous chosen trials during Rule Switch rose earlier in SNAP-treated mice than in controls (Fig. 3h). SNAP-treated mice also reached the 75% rolling-performance threshold at an earlier trial than control mice (Welch’s t test, t(17.31) = 2.41, p = 0.03; Fig. 3i). Together, the session-level and trial-resolved measures show that, under MCHR1 antagonism across both task phases, behavior stabilized earlier after the contingency switch and mice made fewer continued selections of the previously reinforced arm.

### MCHR1 antagonism alters outcome processing and experience-dependent prelimbic responses

To determine whether SNAP altered prelimbic outcome processing, we recorded CaMKIIα-jGCaMP8m signals from prelimbic mPFC during task performance (Fig. 4a). During both Initial Learning and Rule Switch, SNAP enhanced the negative-going mPFC response following correct outcomes, with little effect following incorrect outcomes (Group × Outcome: Initial Learning, β = 1.20, p = 2.31 × 10⁻¹², Fig. 4b,c; Rule Switch, β = 0.93, p = 6.36 × 10⁻⁸, Fig. 4d,e). Because this outcome-dependent effect was already present during Initial Learning, when the measured behavioral indices were unchanged, it could not by itself account for the behavioral difference observed during Rule Switch.

**Figure 4.**
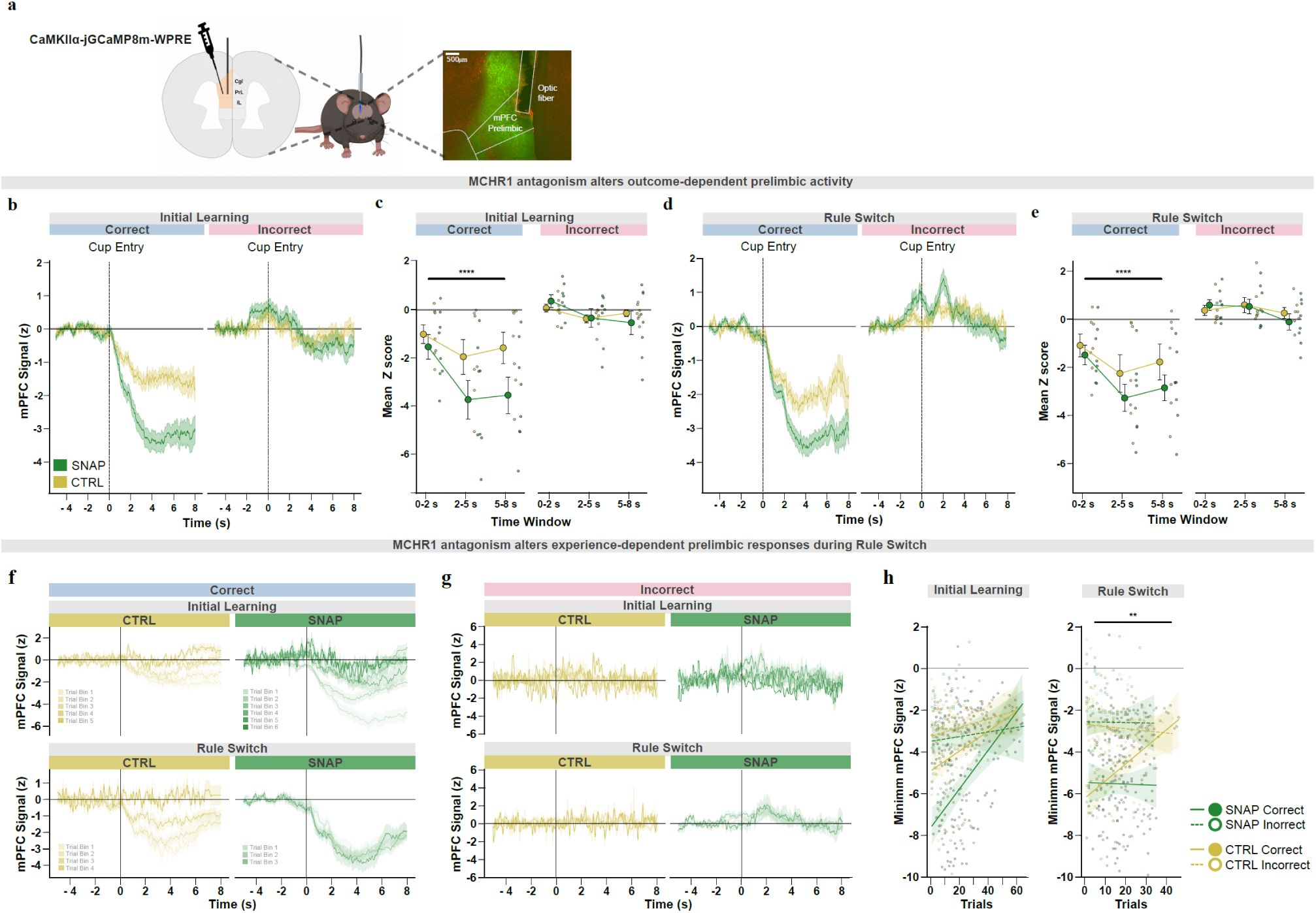
MCHR1 antagonism alters outcome processing and experience-dependent prelimbic responses. (a) Schematic of CaMKIIα-jGCaMP8m expression in prelimbic mPFC, optical-fiber placement, and representative histological verification. (b,d) Cup-entry-aligned mPFC calcium signals during Initial Learning (b) and Rule Switch (d), shown separately for correct and incorrect trials in Control and SNAP-treated mice. (c,e) Mean z-scored mPFC responses during Initial Learning (c) and Rule Switch (e), quantified in early (0–2 s), middle (2–5 s), and late (5–8 s) post-entry windows. (f,g) Correct-trial (f) and incorrect-trial (g) responses across cumulative experience. Trials were ordered continuously across sessions within each phase and grouped into consecutive 12-trial bins independently of outcome; only trials matching the outcome shown were included. Traces show the mean ± s.e.m. across included trials for each group and bin. Bin-specific trial and animal counts are provided in the Source Data. (h) Minimum z-scored mPFC responses in the 5–7 s post-entry window plotted against cumulative trial number, separately by phase, outcome, and treatment group. Trial-level linear mixed-effects models tested Group × Cumulative Trial × Outcome within each phase and Phase × Group × Cumulative Trial × Outcome across phases, with Subject included as a random intercept. All signals were z-scored relative to the −5 to −2 s pre-entry baseline. Control, n = 7 mice (5 male, 2 female); SNAP, n = 8 mice (5 male, 3 female).

We next examined how outcome-related activity evolved with cumulative experience. Correct- and incorrect-trial traces were grouped into consecutive 12-trial bins (Fig. 4f,g), and the minimum response from 5–7 s after cup entry was analyzed across cumulative trials (Fig. 4h). Response trajectories did not differ between groups during Initial Learning. During Rule Switch, the correct-outcome response progressively weakened in Control animals (β = 0.172, SE = 0.055, z = 3.103, p = 0.002), whereas this attenuation was opposed under SNAP (Group × Cumulative Trial: β = −0.219, SE = 0.079, z = −2.772, p = 0.006). This treatment-dependent trajectory differed between correct and incorrect outcomes (Group × Cumulative Trial × Outcome: β = 0.214, SE = 0.096, z = 2.232, p = 0.03).

The combined analysis confirmed that the treatment-dependent trajectory differed between Initial Learning and Rule Switch and depended on trial outcome (Group × Cumulative Trial × Phase: β = −0.230, SE = 0.084, z = −2.728, p = 0.006; Group × Cumulative Trial × Outcome × Phase: β = 0.215, SE = 0.107, z = 2.011, p = 0.04). The critical effect of SNAP was therefore not a general increase in outcome responsiveness, but a change in how the prelimbic response to successful outcomes evolved during Rule Switch. Because each correct choice at this stage both confirms the new contingency and disconfirms the previously rewarded policy, the sustained response under SNAP is consistent with successful feedback remaining influential while the old and new response policies compete.

MCHR1 antagonism did not alter trial-history effects in prelimbic mPFC activity (Supplementary Fig. 2).

## Discussion

The present study extends the functional role of hypothalamic MCH signaling beyond its established involvement in feeding and reward, identifying it as a regulator of the balance between maintaining and revising reward-guided actions. MCHR1 antagonism throughout both task phases did not measurably alter Initial Learning but reduced perseverative responding and accelerated behavioral adaptation after the contingency switch. In parallel, endogenous LH-MCH activity was recruited when contingencies were being established or revised, distinguished rewarded from unrewarded outcomes, and incorporated recent outcome history. At the cortical level, SNAP altered outcome-dependent prelimbic responses in both task phases and, specifically during Rule Switch, prevented the correct-outcome response from weakening as experience accumulated. Together, these findings position MCH signaling at the interface between reward history and behavioral updating, where it may determine how long a previously successful response continues to govern behavior after its consequences change.

Most previous work on LH-MCH neurons has focused on feeding, appetition, reward value, and consummatory behavior ^19–22,35–44^. The present findings reveal a different organizing principle. The same food reward remained available throughout the experiment, yet LH-MCH activity was strongest early in Initial Learning, declined as the action–outcome contingency stabilized, and re-emerged when the rewarded arm was reversed. This profile is inconsistent with a fixed response to food receipt or consumption. Instead, LH-MCH recruitment tracked periods in which the validity of the current action–outcome mapping had to be established or reconsidered.

This learning-stage dependence extends evidence that MCH neurons participate in cognitive processes beyond homeostatic and hedonic control. Natural hypothalamic circuit dynamics contribute to object memorization, REM-active MCH neurons influence the persistence of hippocampus-dependent memories, and MCH signaling contributes to fear extinction and the updating of learned avoidance ^22–24,31^. These findings have sometimes appeared heterogeneous because they encompass memory formation, forgetting, extinction, and behavioral flexibility. A common feature is that each process requires the stability of an existing representation to be regulated according to current demands. The present data extend this framework to appetitive instrumental behavior: LH-MCH recruitment diminished when the current policy remained valid and returned when that policy had to be revised.

Beyond tracking whether a contingency was stable or changing, LH-MCH activity also carried information about recent outcome history. Identical current outcomes elicited different responses depending on whether they followed a recent success or failure. This distinction is behaviorally important because the same rewarded outcome can either confirm an established strategy or support a departure from the immediately preceding unsuccessful response. LH-MCH activity therefore preserved information relevant to evaluating whether ongoing behavior should be maintained or changed.

History sensitivity was evident before as well as after cup entry, although its expression differed between peak and minimum components and between current-correct and current-incorrect trials. This temporal heterogeneity argues against a unitary outcome signal. Instead, the population response appears to combine a state carried into the current choice with activity generated when the outcome becomes available. The supervised analysis provided a complementary trial-wise view of the post-entry peak. Across both phases, model predictions were driven primarily by accumulated performance and task experience, suggesting that LH-MCH responses reflect current outcomes in relation to the animal’s evolving estimate of the action– outcome contingency rather than as isolated reward events. The relative ranking of these experience-related features differed between Initial Learning and Rule Switch, although the phase-specific models and correlated predictors preclude a formal cross-phase comparison.

This history-dependent organization complements recent work showing that frontal cortical populations retain information about previous decisions and outcomes independently of ongoing posture and movement^10^. The present findings show that recent-outcome sensitivity is also evident in a hypothalamic population traditionally associated with feeding and arousal. In contrast, MCHR1 antagonism did not measurably alter trial-history effects in bulk prelimbic activity; its cortical effects were instead expressed in current-outcome processing and in how those responses evolved across experience. This does not imply that prelimbic cortex lacks trial-history information, which may be MCH-independent or confined to neuronal subpopulations not resolved by bulk photometry.

The pharmacological findings reveal a complementary role for MCHR1 signaling in behavioral persistence. SNAP did not measurably alter Initial Learning but reduced repeated choices of the previously rewarded arm and accelerated criterion attainment during Rule Switch. Thus, SNAP did not produce a general improvement across both task phases; it reduced continued expression of the previously rewarded choice after the contingency changed. These findings suggest that MCHR1 signaling contributes to the persistence of previously rewarded actions after their consequences change. Because SNAP was administered throughout both Initial Learning and Rule Switch, the experiment does not isolate an acute effect of MCHR1 blockade during contingency change. The absence of a measurable group difference during Initial Learning argues against a gross acquisition deficit, but does not exclude treatment-dependent differences in the initially acquired representation that subsequently influenced Rule Switch performance.

Renewed LH-MCH activity during Rule Switch may initially appear inconsistent with the facilitation produced by MCHR1 blockade. However, the photometry and pharmacological experiments interrogate different levels of the system. Fiber photometry measures the activity of MCH neurons, which release fast transmitters and other products in addition to MCH peptide, whereas SNAP selectively blocks MCHR1 signaling throughout the brain ^45^. MCH-neuron recruitment can therefore accompany contingency updating without implying that activation of MCHR1 uniformly promotes updating. Similar dissociations have been observed in MCH interactions with accumbens dopamine ^20^. Rather than defining MCH neurons as simply pro- or anti-learning, the present findings indicate that one component of their output—MCHR1 signaling—favors persistence of a previously reinforced response during Rule Switch.

The prelimbic recordings revealed an accompanying change in a cortical system known to integrate outcomes, action history, conflict and task context during adaptive response selection ^1–10,12–18^ . SNAP altered the relationship between correct- and incorrect-outcome responses during both Initial Learning and Rule Switch. Our previous finding that local MCHR1 blockade within prelimbic mPFC facilitates the updating of learned avoidance ^31^ makes a direct cortical contribution plausible. However, because SNAP was administered systemically in the present study, the observed prelimbic effects could also arise indirectly through other MCHR1-expressing circuits, and the present experiment does not distinguish between these possibilities. Because this effect was already present during Initial Learning, when behavior was unchanged, it identifies successful-outcome processing as sensitive to MCHR1 blockade but cannot by itself explain the selective improvement during Rule Switch.

The critical distinction emerged as animals accumulated experience. In Control animals, the correct-outcome response weakened as Rule Switch progressed, whereas it remained pronounced under SNAP. A correct outcome at this stage serves two functions: it confirms the newly rewarded choice and disconfirms the previously successful policy. This pattern raises the possibility that successful outcomes remained influential under SNAP while the old and new response policies competed. However, because SNAP was administered systemically and prelimbic activity was measured rather than manipulated, the present data do not establish that the altered cortical response mediated the reduction in perseverative choice. This interpretation links the altered prelimbic trajectory to reduced perseveration and earlier criterion attainment and accords with evidence that prefrontal populations reorganize when animals abandon outdated rules and establish updated action models^46–49^.

Persistent pursuit of formerly rewarding outcomes despite changed consequences is a feature of compulsive behavior and cognitive inflexibility ^10,50^, whereas MCHR1 signaling has been implicated in food seeking, stress-related behavior, and cocaine reward ^11,51,52^. The present task does not model compulsivity directly. Rather, it isolates persistence of a previously rewarded response after its consequences change, a behavioral process that may contribute to maladaptive rigidity. The findings therefore extend previous work by identifying MCHR1-dependent signaling as a contributor to the inertia of reward-guided response policies. Such stabilization may be adaptive while contingencies remain reliable but maladaptive when prior reward history conflicts with current evidence.

In summary, LH-MCH activity tracked periods of contingency evaluation and reflected current outcomes in the context of recent and accumulated experience, whereas MCHR1 blockade reduced persistence of an outdated response and sustained prelimbic sensitivity to successful feedback during Rule Switch. Together, these findings identify MCH signaling as a regulator of how strongly prior reward history continues to guide behavior when contingencies change, linking hypothalamic neuromodulation to the balance between behavioral persistence and flexibility.

## Acknowledgments

This work was supported by ETH Zurich Grant ETH-24 20-2, awarded to D.P.-R.

## Author contributions

D.P.-R. conceived the study. D.P.-R. and P.K. designed the experiments. P.K. established the experimental setup and performed the experiments; C.C. performed the experiments shown in Fig. 1.; E.B. established the photometry setup and wrote photometry extraction code; P.K., D.B., and D.P.-R. analyzed the data; J.B. performed the supervised regression and SHAP analyses. P.K. prepared the figures. E.B. provided input on figure presentation. D.P.-R. wrote the original draft with input from P.K. and D.B. All authors reviewed and approved the final manuscript.

---

**Supplementary Figure 1.**
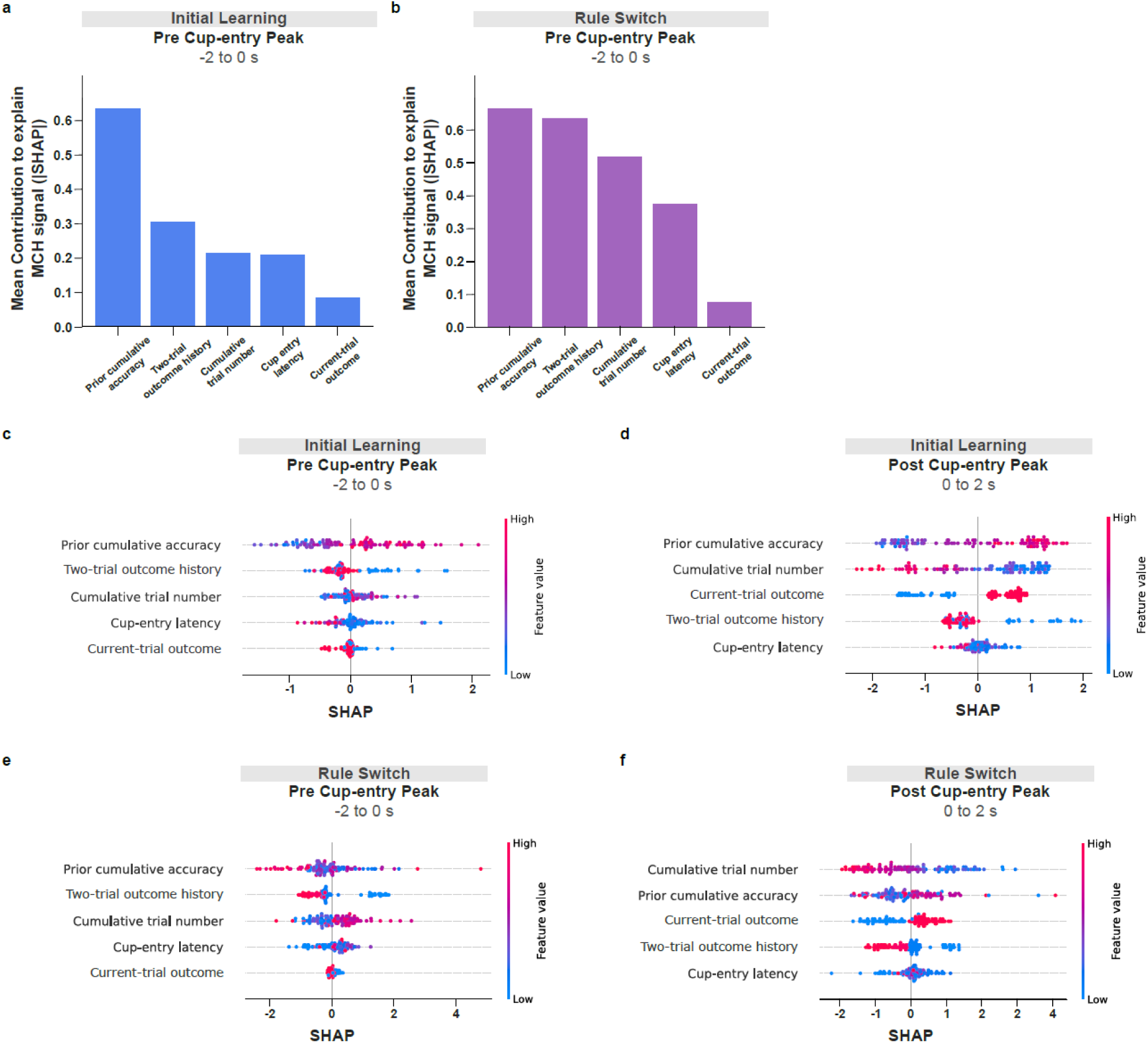
SHAP feature attribution for phase-specific models of LH-MCH peak responses. (a,b) Mean absolute SHAP values for the secondary pre-entry peak models (−2 to 0 s) during Initial Learning (a) and Rule Switch (b). Bars indicate the mean magnitude of each feature’s contribution to model predictions. (c–f) SHAP beeswarm plots for the pre-entry and post-entry peak models during Initial Learning (c,d) and Rule Switch (e,f), respectively. Each point represents one held-out trial. Horizontal position indicates the signed SHAP value, with positive values increasing and negative values decreasing the model-predicted peak. Point color denotes the observed feature value, from low (blue) to high (magenta). Features are ordered by mean absolute SHAP value. For the two-trial outcome-history predictor, point color reflects its numerical encoding (0, 1, 10 and 11, corresponding to the sequences 00, 01, 10 and 11, respectively); SHAP values are interpreted as the contribution of the joint two-trial-history predictor rather than as the effect of a one-unit increase in its numerical value. Mean absolute SHAP values quantify contribution magnitude but not direction. Because SHAP attribution is model-dependent and correlated predictors can share or redistribute importance, attributions are not interpreted as unique, independent or causal effects. Model-performance statistics are provided in Supplementary Table S1, and predictor definitions, coding and formulas are provided in Supplementary Table S2.

**Supplementary Table S1.**
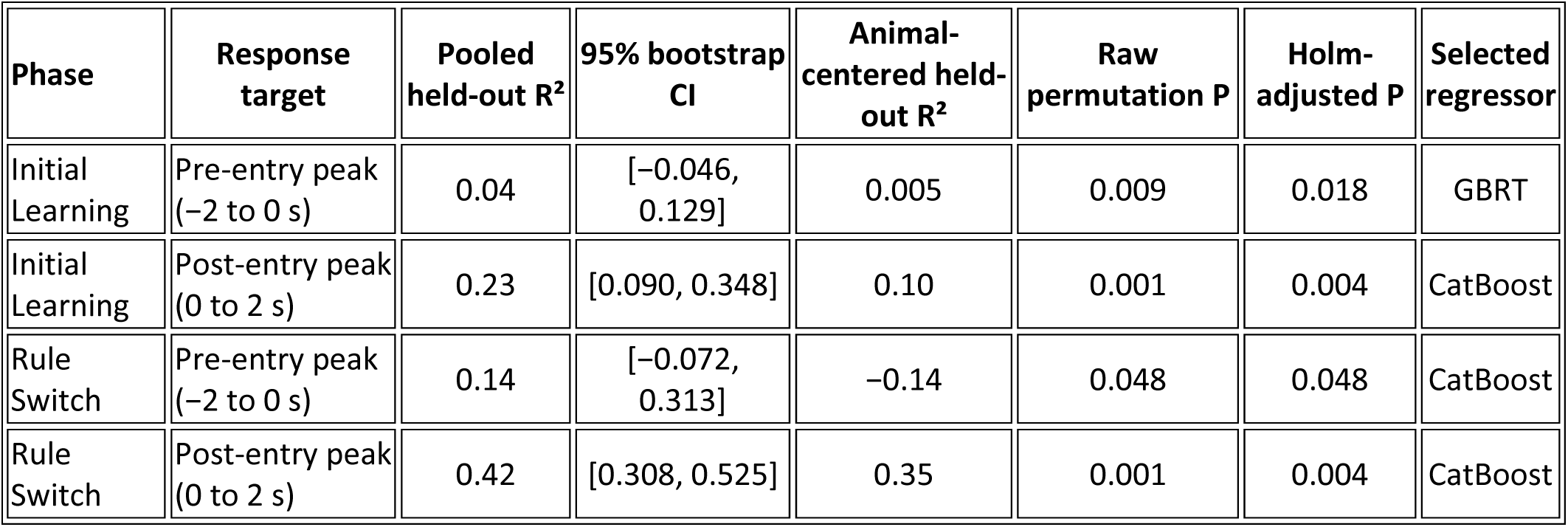
Out-of-sample performance of supervised regression models predicting phase-specific LH-MCH peak responses. Pooled held-out R² quantifies predictive performance on the 30% held-out test trials. Ninety-five percent confidence intervals were obtained using a conditional within-animal trial bootstrap with 2,000 replicates. Raw permutation P values were obtained using fixed-prediction permutation tests in which held-out neural response values were shuffled within animal while model predictions were kept fixed; Holm-adjusted P values correct across the four phase-by-window models. Animal-centered held-out R² is reported as an additional measure of within-animal predictive performance.

| Phase | Response target | Pooled held-out $R^2$ | 95% bootstrap CI | Animal-centered held-out $R^2$ | Raw permutation P | Holm-adjusted P | Selected regressor |
| --- | --- | --- | --- | --- | --- | --- | --- |
| Initial Learning | Pre-entry peak (-2 to 0 s) | 0.04 | [-0.046, 0.129] | 0.005 | 0.009 | 0.018 | GBRT |
| Initial Learning | Post-entry peak (0 to 2 s) | 0.23 | [0.090, 0.348] | 0.10 | 0.001 | 0.004 | CatBoost |
| Rule Switch | Pre-entry peak (-2 to 0 s) | 0.14 | [-0.072, 0.313] | -0.14 | 0.048 | 0.048 | CatBoost |
| Rule Switch | Post-entry peak (0 to 2 s) | 0.42 | [0.308, 0.525] | 0.35 | 0.001 | 0.004 | CatBoost |

**Supplementary Table S2.** Trial-level behavioral predictors used in supervised regression models of LH-MCH peak responses. The same five predictors were used for each phase-by-window model. The table provides the analysis variable name, definition, coding and formula for each predictor. Cumulative trial number and prior cumulative accuracy continued across Initial Learning and Rule Switch without resetting, whereas two-trial outcome history was defined within phase/session.

| Feature name (CSV column) | Definition | Formula |
| --- | --- | --- |
| **Current-trial outcome** (correct) | Binary outcome of the current trial: 1 for a correct/rewarded trial and 0 for an incorrect/unrewarded trial. | $c_t \in \{0, 1\}$ |
| **Two-trial outcome history** (bin2_history) | Joint outcome history of the two immediately preceding completed trials within the same phase/session, ordered as $t-2$ and $t-1$. Each preceding trial was coded as 1 for correct/rewarded and 0 for incorrect/unrewarded. The history resets at each phase/session. Missing early history positions are left-padded with 0, producing the ordered sequences 00, 01, 10, or 11, entered as the numerical values 0, 1, 10, or 11, respectively. | $H_t = 10 \cdot c_{(t-2)} + c_{(t-1)}$<br>(within session) |
| **Cumulative trial number** (global_trial_idx) | One-based cumulative trial number within each animal. It continues across all initial-learning and rule-switch sessions and is not reset at the rule switch. | $G_t = t, t = 1, 2, \dots$<br>(within animal) |
| **Cup-entry latency** (time_to_cup_entry_s) | Time from trial onset to the first genuine outside-to-inside transition into the outcome-appropriate cup zone: the rewarded cup on correct trials and the empty cup on incorrect trials. | $L_t = \min\{\tau_j : z_{(j-1)}=0 \text{ and } z_j=1\}$<br>$z = \text{rewarded if } c_t=1; \text{ empty if } c_t=0$ |
| **Prior cumulative accuracy** (correct_prop_cum_id) | Proportion of all earlier trials that were correct for the same animal. The current trial is excluded, and the calculation continues from initial learning into rule switching without resetting. It is undefined for the first trial. | $A_t = \sum_{i=1}^{(t-1)} c_i / (t-1), t > 1$<br>$A_1 = \text{undefined}$ |

**Supplementary Figure 2.**
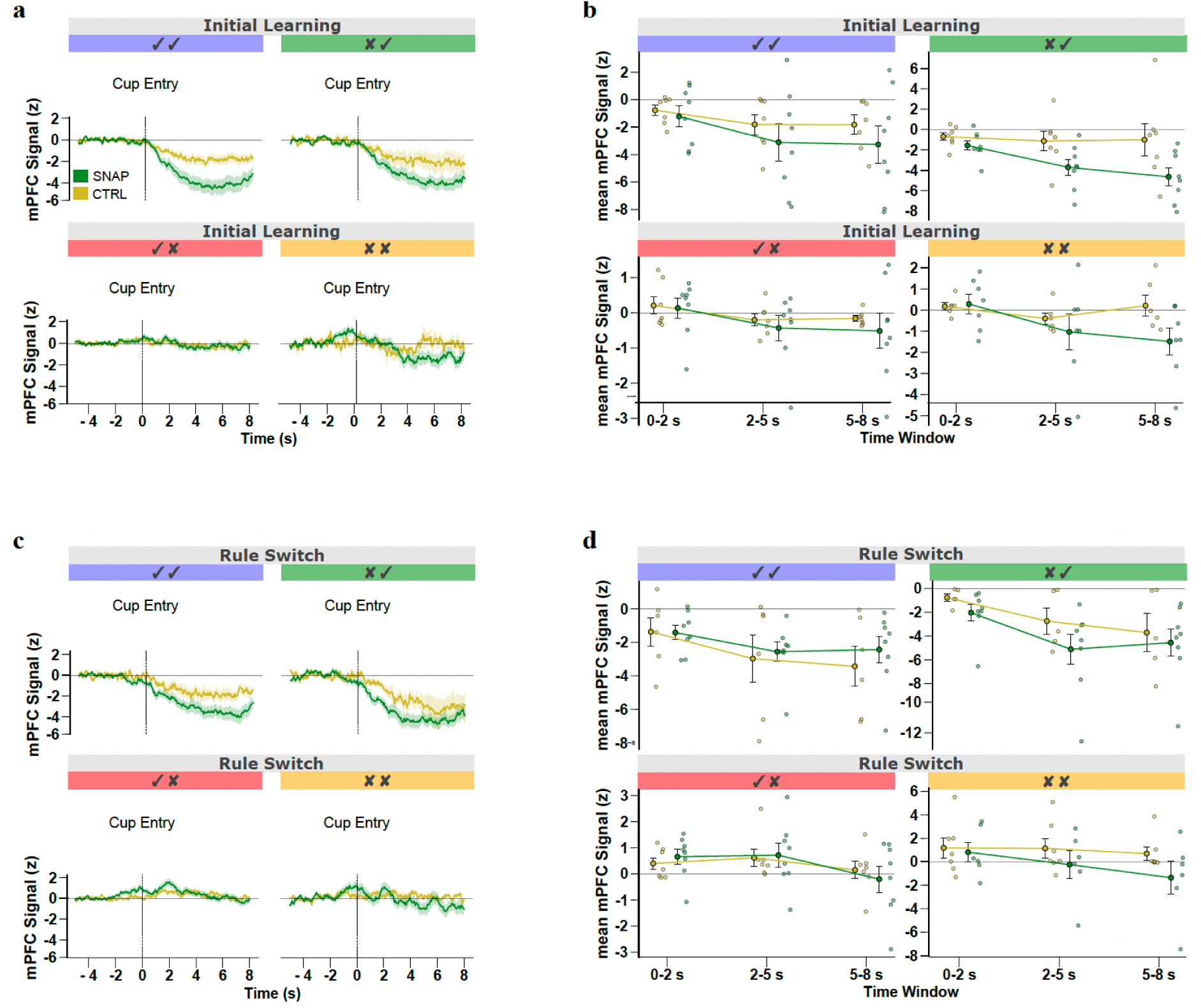
No evidence that MCHR1 antagonism alters trial-history effects in prelimbic mPFC activity. (a,c) Cup-entry-aligned CaMKIIα-jGCaMP8m signals from prelimbic mPFC during Initial Learning (a) and Rule Switch (c), grouped according to the outcome of the immediately preceding trial and the current trial. In each sequence, the first symbol denotes the preceding outcome and the second the current outcome: correct→correct (✓✓), incorrect→correct (✗✓), correct→incorrect (✓✗), and incorrect→incorrect (✗✗). Signals were z-scored relative to the −5 to −2 s pre-entry baseline and are shown as mean ± SEM. (b,d) Mean z-scored responses during Initial Learning (b) and Rule Switch (d), quantified in early (0–2 s), middle (2–5 s), and late (5–8 s) post-entry windows. Trial-level responses were analyzed separately for each phase using linear mixed-effects models including Group, Trial-history sequence, Time Window, and their interactions, with mean pre-entry ΔF/F signal as a covariate and Subject as a random intercept. Planned contrasts compared ✓✓ with ✗✓ trials and ✓✗ with ✗✗ trials and tested whether these history effects differed between Control and SNAP. No significant Group × Trial-history sequence × Time Window interactions were detected during Initial Learning (all *p* ≥ 0.063) or Rule Switch (all *p* ≥ 0.240), providing no evidence that MCHR1 antagonism altered trial-history effects. Trial counts during Initial Learning were: ✓✓, Control *n* = 45, SNAP *n* = 43; ✗✓, Control *n* = 20, SNAP *n* = 34; ✓✗, Control *n* = 42, SNAP *n* = 53; ✗✗, Control *n* = 11, SNAP *n* = 18. During Rule Switch: ✓✓, Control *n* = 21, SNAP *n* = 31; ✗✓, Control *n* = 17, SNAP *n* = 22; ✓✗, Control *n* = 51, SNAP *n* = 49; ✗✗, Control *n* = 31, SNAP *n* = 18. Control, *n* = 7 mice; SNAP, *n* = 8 mice.

## Materials and Methods

### Animals

All procedures involving animals complied with regulations of the Swiss Federal Food Safety and Veterinary Office and were approved by the Cantonal Veterinary Office of Zürich (Animal Welfare Ordinance, SR 455.1). Experiments were performed in adult male and female C57BL/6 mice. Mice were housed under controlled temperature (22 °C) and humidity (55%) on a reversed 12 h light/dark cycle with ad libitum access to water. Behavioral experiments were performed during the dark phase. Before behavioral testing, mice were food deprived overnight. Sex and sample sizes for individual experiments are indicated in the corresponding figure legends.

### Stereotaxic viral injections and fiber implantation

Mice were anaesthetized with isoflurane and placed in a stereotaxic frame (Kopf Instruments). Coordinates are given relative to bregma.

### mPFC photometry

For mPFC fiber photometry experiments, mice received perioperative carprofen (5 mg/kg, s.c.). A small craniotomy was made above prelimbic mPFC, and AAV1-CaMKIIα-jGCaMP8m-WPRE (Addgene; titre ≥ 7 × 10^12 vg/ml) was injected unilaterally using a Hamilton microsyringe (200 nl, undiluted; 50 nl/min) at AP +1.8 mm, ML +0.3 mm and DV −2.0 mm. The needle was left in place for 5 min after infusion to minimize reflux. A fiber-optic implant (200 μm core diameter, 0.48 NA; Thorlabs) was then positioned above the injection site (AP +1.8 mm, ML +0.3 mm, DV −1.9 mm) and secured with dental cement. Mice recovered for at least 3 weeks before behavioral testing.

### LH-MCH photometry

For LH-MCH fiber photometry experiments, mice received meloxicam (Metacam; 2 mg/kg, s.c.) for perioperative analgesia. MCH neurons were targeted using AAV9.pMCH.GCaMP6s.hGH (1.78 × 10^14 gc/ ml; Vigene Bioscience), using the same promoter-based strategy previously validated for selective targeting of MCH neurons in C57BL/6 mice (1-3). Three injections per hemisphere were made into the lateral hypothalamus using a 33-gauge needle mounted on a Hamilton syringe (50 nl per injection, 50 nl/min) at AP −1.35 mm, ML ±0.90 mm and DV −5.70, −5.40 and −5.10 mm. Fiber-optic implants were positioned above the lateral hypothalamus (10° angle; AP −1.35 mm, ML ±1.90 mm, DV −5.0 mm) and fixed to the skull. Mice recovered for at least 10 days before behavioral testing.

### Systemic pharmacology

SNAP-94847 was dissolved in vehicle containing 2% dimethyl sulfoxide (DMSO), (2-hydroxypropyl)-β-cyclodextrin (Merck) and phosphate-buffered saline (PBS). The drug was administered intraperitoneally at 20 mg/kg, with injection volume adjusted to body weight. Vehicle-treated mice received the same injection volume of vehicle alone. Injections were given 45 min before behavioral testing, after which mice were returned to their home cages until the start of the session.

### T-maze behavioral task

Behavior was assessed in a T-maze (Med Associates). Each arm measured 7 × 36.5 cm, and consisted of a start arm, containing an ITI chamber (see fig 1b) and two goal arms, each containing a food cup. The center zone measured 9 × 9 cm and the inter-trial interval (ITI) room measured 7 × 17 cm. The food-cup zone was defined as a 3 cm perimeter around the cup. Animals were video tracked using EthoVision XT 15. Before training, mice were habituated to the maze with empty food cups and to the liquid reward in the home cage.

One goal arm was rewarded with 25 μl strawberry milkshake (Emmi) according to the current task rule, whereas the other cup was empty but carried milkshake odor to minimize odor-guided choices. At the start of each trial, mice were placed in the ITI room for 60s, and then released into the start arm and allowed to choose a goal arm. After a correct choice, mice consumed the reward and were returned to the ITI room. After an incorrect choice, a sliding door confined the mouse in the incorrect arm for 10 s before return to the ITI room. If a mouse did not choose either arm within 60 s, the trial was scored as an omission. The ITI lasted 60 s. The maze was cleaned between trials.

The task consisted of two sequential phases. In the initial learning phase, one arm was consistently rewarded until the mouse reached criterion. In the rule-switch phase, the rewarded arm was switched to the opposite side. Each session consisted of 12 trials. Mice were trained across consecutive sessions until they reached criterion performance, defined as at least 75% correct choices across the session and at least two-thirds correct choices in the final three trials.

### Fiber photometry acquisition and preprocessing

Fiber photometry recordings were acquired using a Doric Lenses system in lock-in mode with simultaneous 405 nm and 465 nm excitation. The 465 nm channel was used to record calcium-dependent fluorescence and the 405 nm channel served as an isosbestic control for motion artefacts and calcium-independent fluctuations.

The 405 nm signal was linearly fitted to the 465 nm signal, and normalized fluorescence changes were calculated as ΔF/F = ((F465 − F405fit) / F405fit) × 100, following previously described approaches (Gunaydin et al., 2014; Lerner et al., 2015). For event-aligned analyses, traces were aligned to cup entry, defined as entry into the 3 cm perimeter surrounding either the rewarded or unrewarded cup. For analyses shown in the main figures, traces were baseline-normalized and converted to z scores using the −5 to −2 s pre-cup-entry window.

### Histology

After completion of the experiments, mice were deeply anaesthetized with pentobarbital and transcardially perfused with PBS followed by 4% paraformaldehyde (PFA). Brains were post-fixed overnight in 4% PFA at 4 °C and cryoprotected overnight in 30% sucrose in PBS. Tissue was frozen on dry ice and sectioned coronally (50 μm for mPFC experiments; 100 μm for LH-MCH experiments). Sections were examined using a fluorescence microscope (Eclipse Ti2, Nikon) to verify virus expression and fiber placement.

### Quantification and statistical analysis

Analyses and data visualization were performed in Python 3.9, and photometry preprocessing was performed in MATLAB R2019b. Data are presented as mean ± s.e.m. Exact sample sizes, experimental units and exact P values are reported in the Results. Unless otherwise stated, tests were two-sided and significance was set at P < 0.05.

#### Behavioral analysis

Behavioral measures shown in Fig. 3 were compared between Control and SNAP-treated mice separately for Initial Learning and Rule Switch phases. Sessions required to reach criterion, total incorrect choices, total correct choices, and the first continuous chosen trial at which rolling performance reached the 75% threshold were compared using two-sided Welch’s t tests. Rolling performance during the Rule Switch phase was calculated across continuous chosen trials using a 12-trial moving window. Carrying the final value forward after an animal completed the task was performed for visualization only and was not used for statistical inference. Omissions were excluded from analyses based on chosen trials but were included in the denominator when calculating session-level percentage correct and determining criterion attainment.

#### mPFC outcome-response analysis

Cup-entry-aligned mPFC responses were quantified from z-scored photometry traces in early (0–2 s), middle (2–5 s), late (5–8 s), and full (0–8 s) post food cup-zone entry windows. Analyses were performed separately for Initial Learning and Rule Switch using trial-level linear mixed-effects models containing Group, Outcome, Time Window, Session, and the specified interaction terms as fixed effects, mean pre-entry ΔF/F signal as a covariate, and Subject as a random intercept. The baseline covariate was defined as the mean ΔF/F signal from −5 to −2 s before cup entry, prior to z-scoring.

Primary models tested whether the correct–incorrect response relationship differed between Control and SNAP using the Group × Outcome interaction. Additional models containing the Group × Outcome × Time Window interaction tested whether this group-dependent outcome effect differed across post-entry intervals. Follow-up contrasts compared Control and SNAP separately within each outcome and time window. The full 0–8 s response was analyzed as an additional summary window.

#### Experience-dependent mPFC analysis

To visualize the evolution of mPFC activity across experience, trials were ordered continuously across sessions within each task phase and assigned to consecutive non-overlapping 12-trial bins, irrespective of trial outcome. Correct and incorrect trials within each bin were then plotted separately. Cup-entry-aligned traces were z-scored relative to the −5 to −2 s pre-entry baseline and averaged across the trials contributing to each treatment group and bin. Shaded regions show s.e.m. across included trials. Because animals reached criterion after different numbers of trials, the number of contributing trials and mice decreased in later bins.

For statistical analysis, the minimum z-scored mPFC response between 5 and 7 s after cup entry was extracted from each trial. Initial Learning and Rule Switch were first analyzed separately using trial-level linear mixed-effects models containing Group, Cumulative Trial, Outcome, and all corresponding interaction terms as fixed effects, with Subject as a random intercept. A combined model additionally included Phase and all corresponding interactions with Group, Cumulative Trial, and Outcome. The Group × Cumulative Trial interaction tested whether SNAP altered the evolution of the mPFC response within each phase, whereas the Group × Cumulative Trial × Outcome interaction tested whether this effect differed between correct and incorrect trials. Phase interactions tested whether these treatment-dependent trajectories differed between Initial Learning and Rule Switch.

#### mPFC trial-history analysis

For the mPFC trial-history analysis shown in Supplementary Fig. 2, trials were classified as correct→correct, incorrect→correct, correct→incorrect, or incorrect→incorrect according to the immediately preceding and current outcomes. Mean z-scored responses were quantified in early (0–2 s), middle (2–5 s), late (5–8 s), and full (0–8 s) post-entry windows. Initial Learning and Rule Switch were analyzed separately using trial-level linear mixed-effects models containing Group, Trial-history sequence, Time Window, and their interactions as fixed effects, with mean pre-entry ΔF/F signal as a covariate and Subject as a random intercept. Planned contrasts compared correct→correct with incorrect→correct trials and correct→incorrect with incorrect→incorrect trials within each treatment group and tested whether these history effects differed between Control and SNAP.

#### LH-MCH photometry analysis

Cup-entry-aligned LH-MCH responses were quantified from z-scored photometry traces. Cup-entry peak responses were defined as the maximum signal from 0 to 2 s after cup entry. Correct- and incorrect-trial peak responses were compared within animals separately during Initial Learning and Rule Switch using two-sided paired Wilcoxon signed-rank tests. Peak responses averaged across phases were compared between correct and incorrect trials using a two-sided paired t test.

To assess learning-related changes, cup-entry peak responses were analyzed separately for Initial Learning and Rule Switch using linear mixed-effects models containing Outcome, Session, and their interaction as fixed effects and Subject as a random intercept. Session was coded as the within-phase session number, beginning with Session 1 in each phase. Models included data from all completed sessions; Session 1, Session 2, and each animal’s final session were selected only for visualization in Fig. 2e. The planned comparison between the final session of Initial Learning and the first session of Rule Switch was performed using a one-sided paired t test.

#### LH-MCH trial-history analyses

For one-back trial-history analyses, trials were classified according to the immediately preceding and current outcomes: correct→correct, incorrect→correct, correct→incorrect, and incorrect→incorrect. Peak and minimum values were calculated separately in the pre-entry window from −2 to 0 s and the post-entry window from 0 to 2 s relative to cup entry. Within current-correct trials, responses following a correct versus incorrect preceding trial were compared within animals. The corresponding comparison was performed within current-incorrect trials. Two-sided paired tests were used, and p values were corrected for multiple comparisons using the false-discovery-rate procedure.

#### Supervised prediction of MCH response measures and SHAP analysis

Because multiple behavioral variables co-occurred within individual trials, supervised regression models were used to determine which behavioral features were informative about trial-to-trial variation in the MCH signal. Separate models were trained to predict four neural response measures defined by task phase and temporal window: the pre-entry peak during Initial Learning, the post-entry peak during Initial Learning, the pre-entry peak during Rule Switch, and the post-entry peak during Rule Switch. Pre-entry values were calculated from−2 to 0 s relative to cup entry, and post-entry values from 0 to 2 s. Cup entry was defined as the first entry into the rewarded cup zone on correct trials or the empty-cup zone on incorrect trials. Because the supervised analysis was designed as a multivariable extension of the primary cup-entry peak analysis, peak responses were used as prediction targets, whereas minimum responses were retained for the targeted trial-history analysis.

The preceding event-related analyses used traces baseline-corrected to the −5 to −2 s interval. In contrast, the regression targets were extrema extracted from the ΔF/F signal across the complete recording, without additional event-specific baseline subtraction. The peak was defined as the maximum time-weighted mean of the ΔF/F signal within a 0.5-s window rather than as a single maximum sample. Before calculating these interval means, isolated one-sample artifacts were identified using a time-based local Hampel filter and replaced with the corresponding local median. Longer sequences of elevated samples were retained to avoid removing potentially genuine biological transients.

Trial-level behavioral features were merged by animal ID, task phase, and trial. The same five behavioral features were used for Initial Learning and Rule Switch: current-trial outcome, two-trial outcome history cumulative trial number, cup-entry latency, and prior cumulative accuracy. Current-trial outcome was coded as 1 for a correct trial and 0 for an incorrect trial. Two-trial outcome history were represented by a single numerically encoded feature ordered as t−2 and t−1. Values of 0, 1, 10, and 11 represented the sequences 00, 01, 10, and 11, respectively. The history reset at each phase/session; when fewer than two preceding outcomes were available, missing leading position(s) were padded with 0. Because the selected tree-based regressors used this feature in threshold-based splits, the numerical encoding did not impose a linear proportional relationship between code distance and the predicted MCH response. However, the encoding imposed an ordering of the four history states; SHAP attribution was therefore interpreted as the contribution of the joint two-trial-history feature, rather than as the effect of a one-unit increase in its numerical value. Cumulative trial number was defined within each animal and continued across Initial Learning and Rule Switch without resetting at the rule switch. Cup-entry latency was defined as the time from trial onset to the first genuine outside-to-inside transition into the outcome-appropriate cup zone: the rewarded cup on correct trials and the empty cup on incorrect trials. Prior cumulative accuracy was the proportion of all preceding trials that were correct for the same animal; the current trial was excluded and the calculation continued from Initial Learning into Rule Switch without resetting. Complete feature definitions, coding, and formulas are provided in Supplementary Table S2.

For each of the four phase-by-window targets, Initial Learning and Rule Switch were maintained as separate datasets. Within each animal, trials were divided into a training subset used for model selection and hyperparameter optimization (70%) and a held-out test subset (30%). The two subsets contained non-overlapping trials. The held-out subset was not used for model fitting or tuning and was subsequently used for final performance evaluation and SHAP analysis. The regression model family and hyperparameters were selected within the training subset using Optuna by maximizing the mean validation R^2^ across five trial-wise cross-validation folds constructed within animal. After model selection, the final tuned regressor was refitted using the complete training subset and used to predict the held-out test observations. Final predictive performance was quantified as pooled R^2^ on the held-out test subset. The selected model family differed for one of the four targets: GBRT was selected for the Initial Learning pre-entry peak, whereas CatBoost was selected for the Initial Learning post-entry peak and for both Rule Switch peak models.

In addition to standard pooled R², an animal-centered R² was calculated to quantify trial-to-trial predictive performance after removing stable differences in mean signal level between animals. For each animal, the observed responses and corresponding model predictions in the held-out test set were separately centered by subtracting their respective animal-specific test-set means. R² was then calculated after pooling all centered held-out trials across animals:

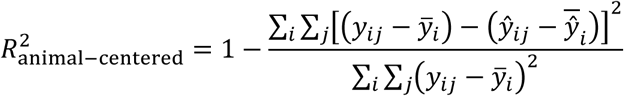

Here, *y_ij_* and 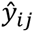 denote the observed and predicted responses for held-out trial (j) from animal (i), and 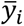 and 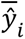 denote the corresponding animal-specific means across held-out trials. This centering was applied only when calculating the performance metric and did not alter model fitting or the original predictions. The resulting statistic is a pooled, trial-weighted measure and not the mean of separately calculated animal-specific R² values.

Uncertainty in held-out model performance was quantified using a conditional within-animal trial bootstrap. In each of 2,000 bootstrap replicates, held-out trials were sampled with replacement separately within each animal while retaining the original number of test trials contributed by that animal. Observed responses and their corresponding fixed model predictions were resampled together, pooled held-out R^2^ was recalculated, and percentile-based 95% confidence intervals were obtained from the resulting bootstrap distribution. These intervals therefore quantify trial-sampling uncertainty conditional on the fitted model, the selected train–test split, and the recorded animals.

To test whether the correspondence between observed and predicted responses in the held-out trials exceeded that expected by chance, we performed a fixed-prediction permutation test. For each target, after model selection and fitting had been completed using the training data, the fitted model’s held-out predictions were kept fixed, and the held-out neural response values were shuffled separately within each animal 999 times. This generated a null distribution of pooled held-out R^2^ values. The one-sided empirical permutation P value was calculated as 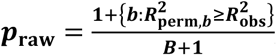, where B=999 valid permutations. The resulting P values were corrected across the four modeled peak targets using the Holm procedure.

For completeness, the same model-fitting and performance-evaluation procedure was applied to the two pre-entry peak measures. Model-performance statistics for these analyses are reported in Supplementary Table S1, and the corresponding SHAP profiles are shown in Supplementary Fig. 1.

The observed and resampling-derived model-performance statistics for all four models are reported in Supplementary Table S1. The training and held-out partitions contained non-overlapping trials, although each animal contributed trials to both partitions. These results therefore establish generalization to held-out trials from the recorded animals, but not to animals absent during training.

Contributions of individual behavioral features to model predictions were quantified using SHapley Additive exPlanations (SHAP). SHAP values were calculated for held-out test observations using the final model fitted exclusively on the training data. For each feature, the mean absolute SHAP value was calculated across held-out observations and used to quantify the magnitude of its contribution to model predictions. Mean absolute SHAP values were used for the feature rankings shown in Fig. 2m,n, whereas the distributions and directions of individual feature contributions are shown in the corresponding SHAP beeswarm plots in Supplementary Fig. 1. In each beeswarm plot, every point represents one evaluated held-out trial; its horizontal position indicates whether the feature shifted the predicted MCH peak upward or downward, while color represents the corresponding feature value.

Mean absolute SHAP values quantify the magnitude, but not the direction, of each feature’s contribution. Because SHAP attribution is model-dependent and correlated features can share or redistribute importance, these rankings describe how the fitted models used the available features rather than unique, independent, or causal effects of individual behavioral features on MCH activity.

#### Sex as a biological variable

Male and female mice were initially inspected separately for sex-dependent effects. Because no significant sex differences were detected in the behavioral datasets analyzed here, data from both sexes were pooled within experimental groups for the final analyses.

